# Improving Deconvolution Methods in Biology through Open Innovation Competitions: An Application to the Connectivity Map

**DOI:** 10.1101/2020.01.10.897363

**Authors:** Andrea Blasco, Ted Natoli, Michael G. Endres, Rinat A. Sergeev, Steven Randazzo, Jin H. Paik, N. J. Maximilian Macaluso, Rajiv Narayan, Xiaodong Lu, David Peck, Karim R. Lakhani, Aravind Subramanian

## Abstract

Do machine learning methods improve standard deconvolution techniques for gene expression data? This paper uses a unique new dataset combined with an open innovation competition to evaluate a wide range of gene-expression deconvolution approaches developed by 294 competitors from 20 countries. The objective of the competition was to separate the expression of individual genes from composite measures of gene pairs. Outcomes were evaluated using direct measurements of single genes from the same samples. Results indicate that the winning algorithm based on random forest regression outperformed the other methods in terms of accuracy and reproducibility. More traditional gaussian-mixture methods performed well and tended to be faster. The best deep learning approach yielded outcomes slightly inferior to the above methods. We anticipate researchers in the field will find the dataset and algorithms developed in this study to be a powerful research tool for benchmarking their deconvolution methods and a useful resource for multiple applications.

## 1 Introduction

A recurring problem in biomedical research is how to isolate signals of distinct populations (cell types, tissues, and genes) from composite measures obtained by a single analyte or sensor. This deconvolution problem often stems from the prohibitive cost of profiling each population separately^1, 2^ and has important implications for the analysis of transcriptional data in mixed samples^3–6^, single-cell data^7^, the study of cell dynamics^8^, and imaging data^9^. In the context of gene expression analysis, available deconvolution approaches offer several advantages but also have various limitations^3, 10^. Machine learning approaches present a promising route towards improvement but implementing and benchmarking these methods can be challenging. A prominent issue is the scarcity of realistic “ground truth” data for training and validation. Existing methods are often trained, and have their results validated, on synthetic data rather than biologically mixed expression data, which are more difficult to generate and evaluate. But even when ground truth data is available, other difficulties arise that may prevent an effective evaluation of machine learning approaches, such as complex parameter optimization which requires substantial experience in more advanced machine learning techniques.

In this paper, we addressed these challenges using an open innovation competition. The goal of the competition was to solve a gene-expression deconvolution problem with application to the Connectivity Map (CMap), a catalog of over 1.3 million human gene-expression profiles^1^. To reach such a massive scale, CMap focuses on a reduced representation of the transcriptome, consisting of approximately 1,000 human genes, called “landmarks.” These genes were selected to capture a large portion of the cell’s transcriptional state, thus yielding significant cost reductions compared to traditional methods (RNA-sequencing). However, the data-generation technology (a bead-based multiplex assay called L1000) is limited to 500 different bead colors per sample. Hence, to measure 1,000 genes per sample, CMap coupled each of the available bead types with a gene pair, using a k-means algorithm called “dpeak” to separate the expression of 1,000 genes from the 500 bead measurements.

The goal of the competition was to develop methods to improve upon the dpeak algorithm. Examples from past competitions have shown that machine learning algorithms developed through open competitions tend to outperform solutions derived through conventional means^11–13^. Accordingly, we wanted to see if machine learning solutions developed through the contest could yield considerable improvements over the current dpeak algorithm. We generated a novel dataset for the contest, which is now available online (S1 Data).

The dataset consists of a collection of gene-expression profiles for 122 different perturbagens, both short hairpin RNA (shRNA) and compound treatments at multiple replicates, for a total of over 2,200 gene expression experiments. The same set of experiments was profiled twice while varying the detection mode for acquiring the data. Detection varied between a dual procedure (DUO), two genes per bead barcode, and a uni procedure (UNI), one gene per bead barcode. The UNI data served as the ground truth against which deconvolution procedures applied to the DUO data were compared.

The contest lasted 21 days and was run on Topcoder (Wipro, India), a popular crowd-sourcing platform. Competitors had access to both UNI and DUO datasets, which could be used to inform the development of their deconvolution solutions. Performance evaluation was based on pre-specified metrics of accuracy and computational speed. A prize purse of $23,000 in cash was offered to competitors as an incentive to be divided among the top 9 submissions. To align incentives and prevent problems like overfitting, the cash prizes were awarded based on the performance evaluation on the *holdout* (not seen by the competitors during the competition) subset of the data.

## 2 Methods

Figure 1 shows a schematic illustration of the contest’s main features including a problem statement, the data, and the scoring function used to evaluate submissions.

**Figure 1.**
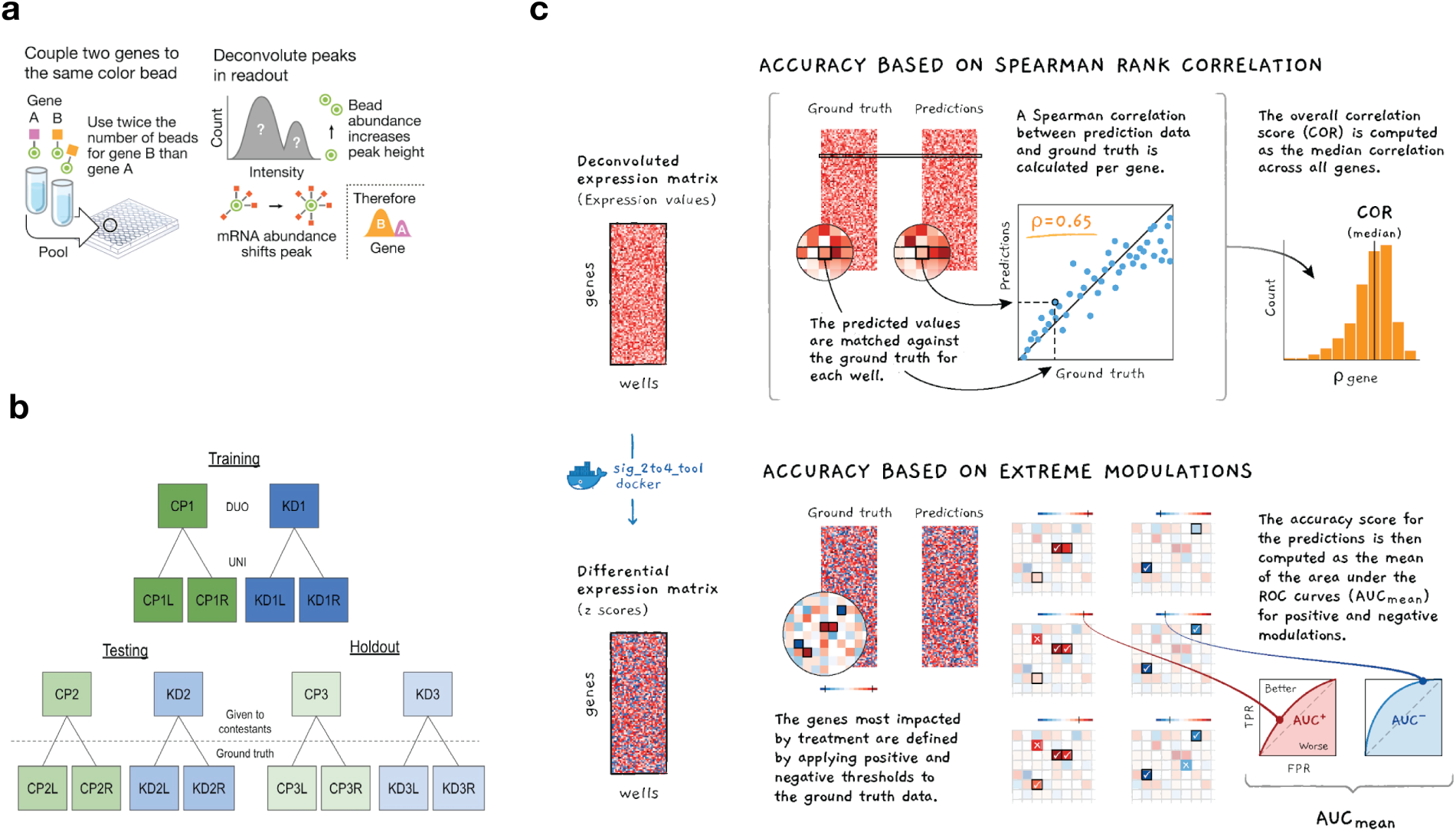
Schematic illustrating the computational problem, generated data, and scoring function. Panel (a) provides a schematic description of the L1000 DUO detection mode and the associated deconvolution problem to be addressed by the contestants. Panel (b) shows the data generated for the contest comprising 6 different sets of perturbational experiments with 3 plates of compound (CP) and shRNA treatments (KD) each. Each plate was detected using DUO (2 genes per analyte) and UNI (one gene per analyte) with UNI serving as the ground truth. Contestants were given two plates of data for training their models offline; a second set of two plates was used during the contest for testing and to populate the live leaderboard; and the third set of two was used as holdout to determine the final contestant placements. Panel (c) illustrates the accuracy component of the scoring function that was used to evaluate the solutions submitted by the competitors. A solution’s overall accuracy score was the product of the genewise Spearman rank correlations with ground truth (DECONV data) and the AUC of extreme modulations (DE data).

The contest’s problem statement described the deconvolution task and the current solution in detail. The key insight was that the typical distribution of measurements for each gene pair has two peaks, the size of which should reflect the relative proportion of the genes in the sample. L1000 takes full advantage of this statistical property by: (1) pairing genes optimally, trying to maximize the average difference in their expression levels, and (2) mixing genes in a 2:1 proportion, which enables the correct assignment of peaks to each gene within the pair (see Subramanian et al. ^1^ for details). Then, it uses a deconvolution approach to partition the composite measurements for each profile into *k* clusters (by minimizing the within-cluster sum of squares) and associate the largest (smallest) cluster to the gene with higher (lower) bead proportion, assigning the cluster’s median value to the corresponding gene.

The datasets consisted of six 384-well perturbagen plates, each containing mutually exclusive sets of compound and shRNA treatments (S1 Table and S2 Table show a complete list). Multiple cell lines and perturbagen were used to avoid any potential over-fitting. The compound and shRNA plates were arbitrarily grouped into pairs, and to avoid any potential ‘information leakage’ each pair was profiled in a different cell line. The resulting lysates were amplified by Ligation Mediated Amplification (LMA, Subramanian et al. ^1^). The amplicon was then split and detected in both UNI and DUO detection modes, resulting in three pairs of data generated under comparable circumstances. The training data was available for all the contestants to develop and validate their solutions offline. The testing data was used for submission evaluation during the contest and to populate a live leaderboard. The holdout data was used for final evaluation, thus guarding against over-fitting. Prizes were awarded based on performance on the holdout dataset.

The scoring function combined measures of accuracy and computational speed (S1 Appendix). The accuracy metric was the product of two different metrics. The first was the average genewise Spearman’s rank correlation between the deconvoluted expression values and the ground truth. The second was the Area Under the receiver operating characteristic Curve (AUC) in the prediction of extremely modulated genes. Speed was measured by executing each submission on comparable multi-core machines, thus allowing competitors to employ multithreading techniques, and the corresponding score was the average runtime in units of the benchmark runtime.

## 3 Results

The contest attracted 294 participants who made 820 submissions using a variety of different methods (S1 Table).

### 3.1 Top four ranked approaches

The winning solution (by a competitor from the United States with a degree in Physics from the University of Kansas) used a random forest algorithm. The algorithm combined predictions from 10 different trees trained on 60 derived data features. These features included a combination of low-peak and high-peak estimates for each gene pair and aggregate measures that are sensitive to systematic bias at the perturbagen, analyte, and plate level.

The second solution (by a competitor from Poland with a Master’s degree in Computer Science from the Lodz University of Technology) used the Expectation-Maximization (EM) algorithm to fit a mixture of two log-normal models to the data for each gene pair. Instead of assuming a priori probability (the 2:1 ratio) of assignment to clusters, the algorithm learned it from the data by fitting a plate-wide distribution of cluster sizes.

The third solution (by a competitor from India with a bachelor’s degree in Computer Science) was a fast k-means algorithm with a random initialization procedure that tends to avoid local minima and is more robust to extreme outliers.

The fourth solution (by a competitor from Ukraine with a bachelor’s degree in Computer Science from the Cherkasy National University) used a Convolutional Neural Network (CNN). This algorithm first filters and transforms the data into a 32-bin histogram for each pair of genes. Then, it uses the U-net architecture^14^ to provide an adequate representation of the data. Then, it assigns each of the 32 bins to one of the two genes for each pair and predicts the median value. This final step uses two subnetworks with the same architecture with a mean squared error loss function (instead of the Spearman correlation used for scoring).

For each method, we generated the deconvolution data (DECONV) and the corresponding differential expression (DE) values as in^1^. Performance was then evaluated on the hold-out data.

### 3.2 Clustering by method and perturbagen type

First, we visually examined the quality of solutions using a two-dimensional t-SNE projection15, which was run once on each of the entire DECONV and DE datasets.

The DECONV data clustered well by perturbagen type and less by algorithm type (Figure 2), although with some notable commonalities in the predictions generated by similar approaches (Figure 2, **d**). For example, the decision tree regressor (DTR) algorithms had similar ‘footprints’ in the projection, as do the k-means and Gaussian mixture model (GMM) algorithms.

**Figure 2.**
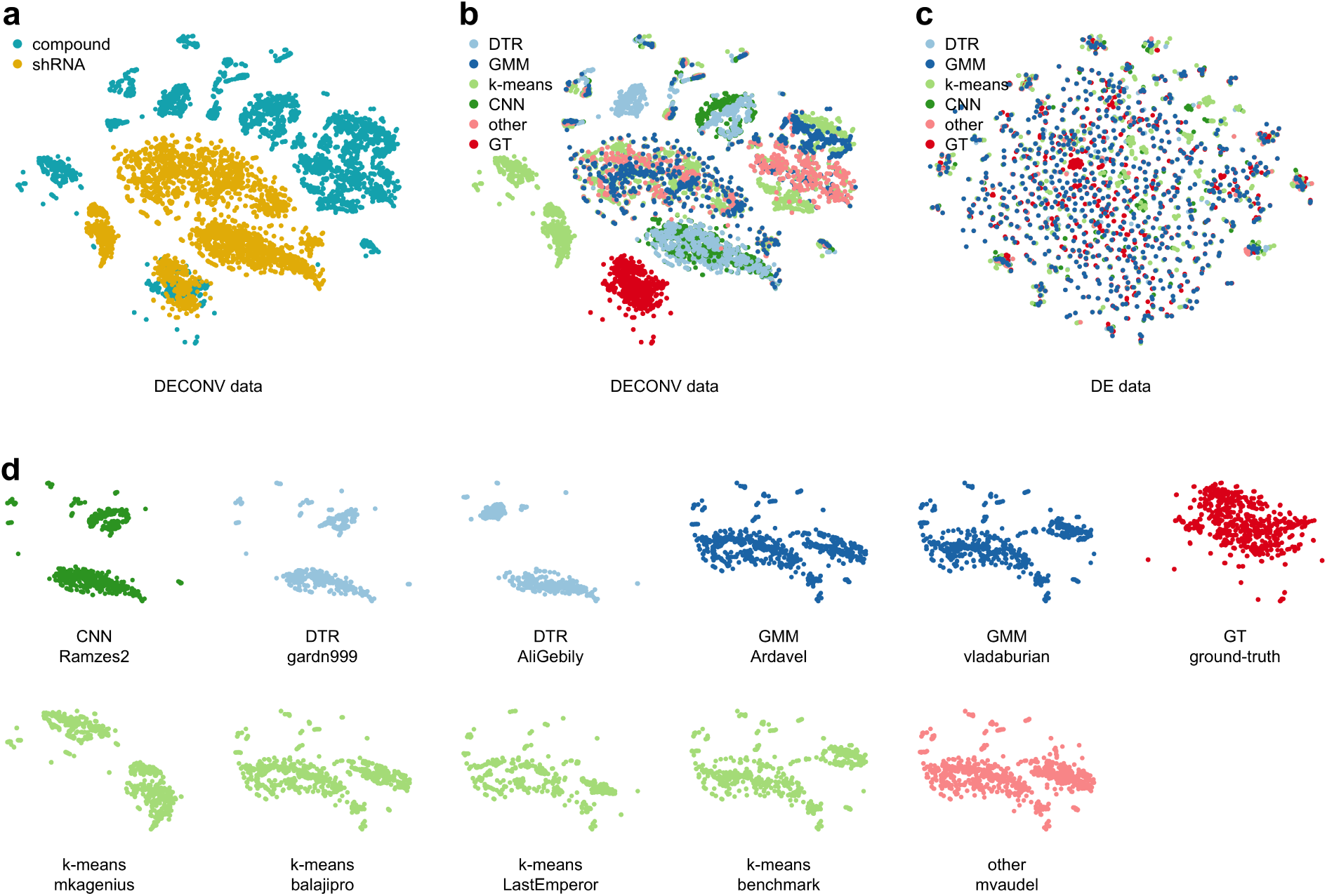
Clustering of solutions. Plot of two-dimensional t-SNE projection with all the samples generated by UNI ground truth (GT) and by applying a deconvolution algorithm to DUO data. t-SNE was run once on all deconvolution data (DECONV) and once on all differential-expression (DE) data that include the 2 plates of holdout data, one each containing compound and shRNA treatments. The resulting projections were colored and subset to generate the following panels: **a**, DECONV data colored by perturbagen type; **b**, DECONV data colored by algorithm type; **c**, DE data colored by algorithm type; and **d**, DECONV data colored by algorithm type and stratified by each implementation.

After the transformation to DE data, however, the t-SNE projection was more homogenous, with no particular clustering by perturbagen and algorithm type (Figure 2, **c**). The lack of clustering in the DE data was reassuring given standard CMap analysis is performed on DE values.

### 3.3 Deconvolution accuracy

Then, we assessed the deconvolution accuracy of each approach by looking at the genewise Spearman rank correlation between the UNI and DUO values.

Based on the analysis of the benchmark, we expected: 1) a higher deconvolution accuracy in samples from shRNA experiments than in those from compounds (Figure 3, a), and 2) a higher deconvolution accuracy for genes in high bead-proportion relative to those in low bead-proportion (Figure 3, b). Intuitively, the deconvolution accuracy should be higher in more focused experiments and for genes with a higher number of beads.

**Figure 3.**
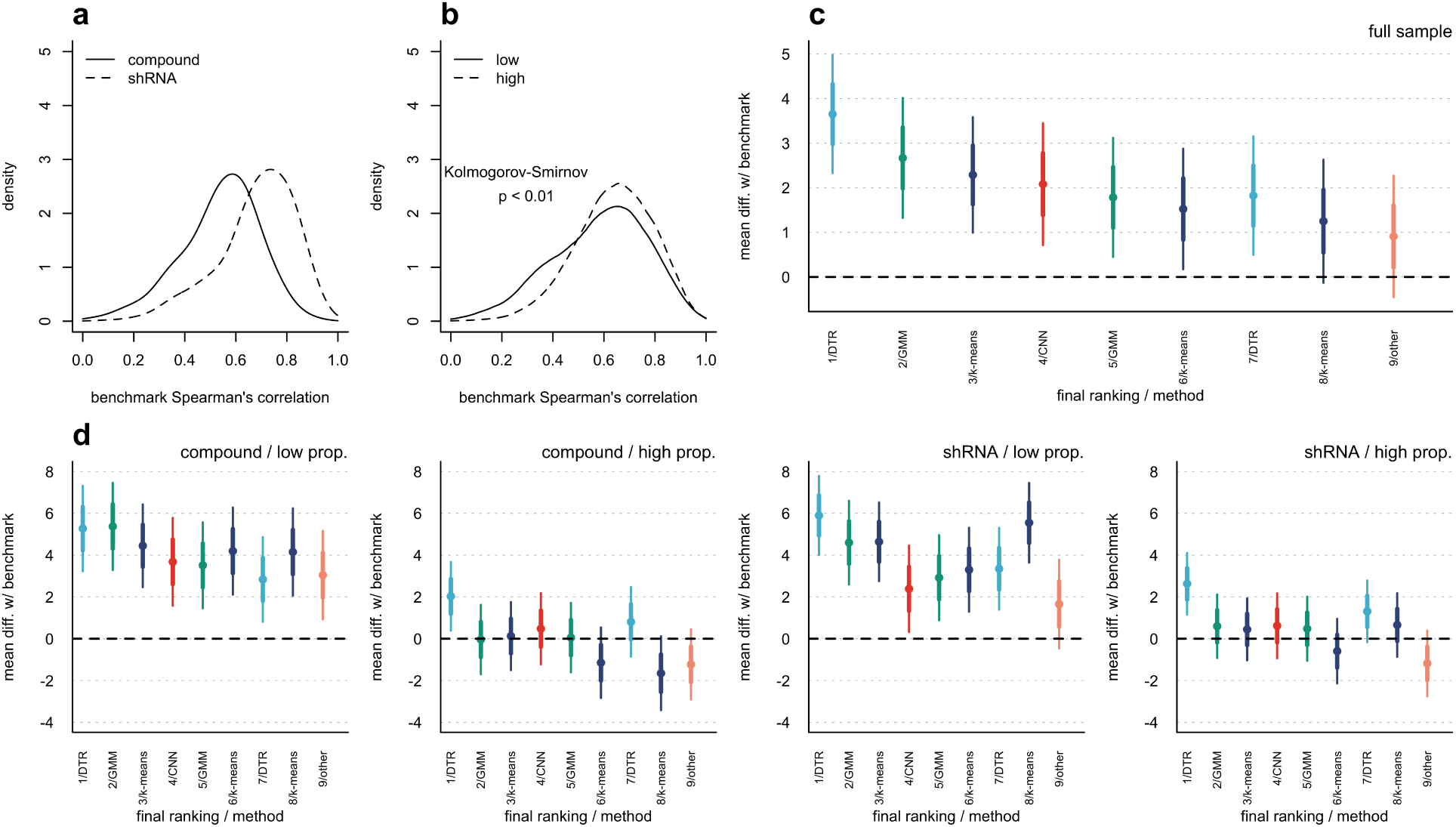
Correlation between ground-truth and deconvolution samples. **a, b** show the empirical distributions of the median correlation between the ground-truth and the deconvolution data for the holdout plates. The median correlation was computed across all the submissions, including the benchmark. **a** shows the distributions for the samples of shRNA and compound treatments. **b** shows the distributions for the subset of genes in low and high bead proportion. **c, d** show the gene-wise mean and confidence intervals (95%) for the correlation coefficient for each of the top-nine performing solutions and the benchmark (BM). **c** shows the gene-wise mean correlation by implementation for the full sample. **d** shows the gene-wise mean correlation by implementation stratified by perturbagen type and bead proportion.

On average across all samples, eight out of the top nine ranked approaches (90%) were significantly better than the benchmark (Figure 3, c). However, at a disaggregated level, improvements were significant for the 488 genes in low bead-proportion and not significant for those in high bead-proportion (Figure 3, d-g). Only the winning method achieved significant improvements in accuracy for genes in both high and low bead proportions.

To evaluate the extent to which the winning algorithm outperformed the others, we ranked the top-nine algorithms by the average correlation metric for each gene (1 = highest, 9 = lowest). We then computed the percentage of genes for which a given algorithm was ranked first. The winner was ranked first for 30% of the genes, followed at some distance by the second-placed gaussian-mixture method (20%), and by the CNN method (13%). Thus, the top two submissions combined outperformed the rest for about half of the genes. Even so, all but a few algorithms were the best performers for at least 5% of the genes, suggesting some complementarity between these algorithms.

### 3.4 Detection of extreme modulations

To evaluate the accuracy at the DE level, we used the detection accuracy of extreme modulations (genes notably up- or down-regulated by perturbation). We used the UNI data with DE values above a threshold as the ground truth; and we evaluated the detection accuracy of each solution by computing the corresponding AUC for each perturbagen type.

The detection accuracy of extreme modulations was generally high for both shRNA and compound samples (AUC > 0.87 and AUC > 0.91, respectively), with the competitors achieving notable improvements over the benchmark (Figure 4, a). Compared to the benchmark, the winning solution detected about 4,000 less extreme modulations (40,800 and 44,200, respectively), thus being more conservative. However, when we restricted the comparison to extreme modulations detected by UNI as well (thus controlling for detection precision), the winning solution detects about 1,500 more extreme modulations than the benchmark (27,100 and 25,600, respectively), representing a sensible 6% increase in “true” detections.^1^

**Figure 4.**
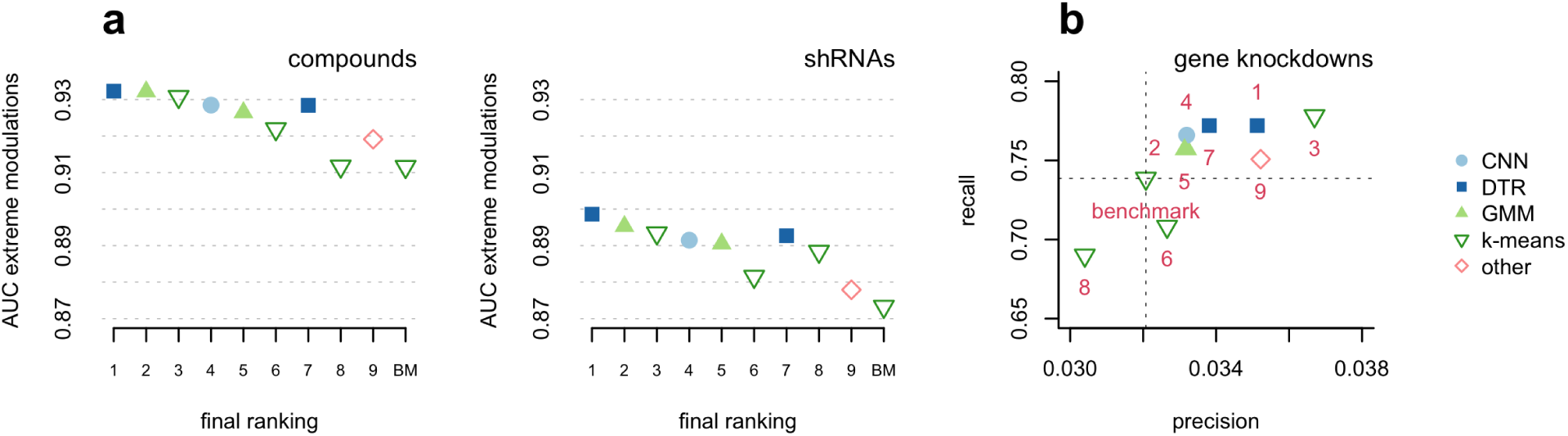
Detection of extreme modulations and targeted knockdown genes. This figure shows the AUC for the predicted differential expressions (DE data) obtained by the top-nine performing methods and the k-means benchmark (dpeak) and the corresponding ground-truth extreme modulations (as detected in the UNI data) both for the shRNA and compound samples (**a**). It also shows the computed recall and precision of the top-nine methods and the benchmark for the detection of the targeted knockdown genes for a subset of shRNA experiments (**b**).

We complemented the above analysis by using targeted gene knockdown (KD) experiments as the ground truth for a subset of data. These are experiments in which a landmark gene was targeted by an shRNA, and hence we expect to observe a significant decrease in expression for the targeted gene. We evaluated the KD detection accuracy of each solution by computing the corresponding percentage of successful KD genes identified or recalled by the algorithm (defining a successful KD as one gene in which the DE value and the corresponding gene-wise rank in the experiment are less than a given threshold, −2 and 10 respectively). We computed the percentage recall for the UNI data as well, which yielded an estimate of the maximum achievable recall of 0.80. Relative to this level, nearly all algorithms achieved a good recall and precision, with values that were higher than the benchmark solution for all but two methods (Figure 4, b).

### 3.5 Reduced variation across replicate samples

To evaluate the reproducibility of the results, we leveraged the several replicate samples for each shRNA and compound experiment in our dataset (about 4 and 10 replicates, respectively). We computed the mean gene-wise coefficient of variation (CV) for each method, which is a measure of inter-replicate variability computed as the average ratio between the interquartile range and the median value across all the replicates. Using this measure, we found all solutions achieved significant improvements over the benchmark (Figure 5); and the winning method, which was the most accurate on average, also achieved the lowest inter-replicate variation overall.

**Figure 5.**
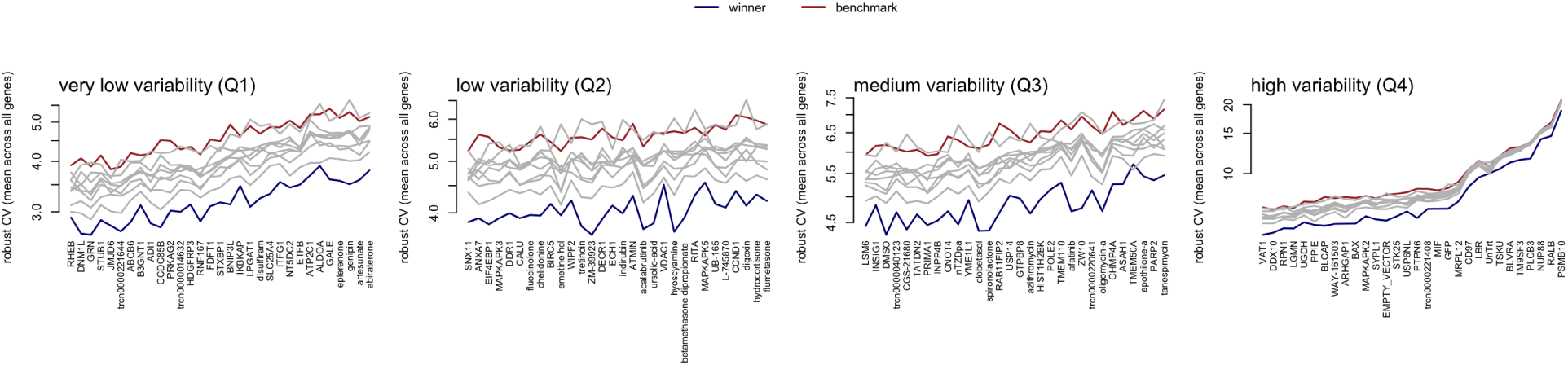
Variation across replicates for the top-four performing methods and the (dpeak) benchmark. This figure shows the mean coefficient of variation for each compound perturbagen in our sample. The mean coefficient of variation is computed as the average of the ratio between the interquartile-range and the median of the deconvoluted values across 10 replicates. The perturbagens on the x-axis are ordered by increasing mean coefficient of variation of the benchmark and split into separate panels by quantiles to make differences more visible.

### 3.6 Computational speed

The speed improvements over the benchmark were substantial. While dpeak took about 4-5 minutes per plate, the fastest algorithm took as little as 5 seconds per plate (more than a 60x speedup compared to the benchmark) and the slowest was well below one minute. These speed improvements are not directly attributable to the use of multiple cores, since both the benchmark and contestant algorithms leverage multi-core techniques. We observed no particular trade-off between speed and accuracy.

### 3.7 Ensembles

Lastly, we assessed the performance of ensembles combining the predictions of different computational methods by taking the median value across all 10 predictions (including the benchmark). By focusing on the subset of the data with shRNA experiments (ignoring the data with compound experiments), the performance in both Spearman correlation and the AUC metrics of the ensemble tended to increase with the number of models involved (Figure 6). However, the maximum performance in both metrics tended to plateau (or even decrease) after combining the results of 3 or more models. This result suggested limited gains from having ensembles, although it may be worth exploring more sophisticated aggregation approaches.

**Figure 6.**
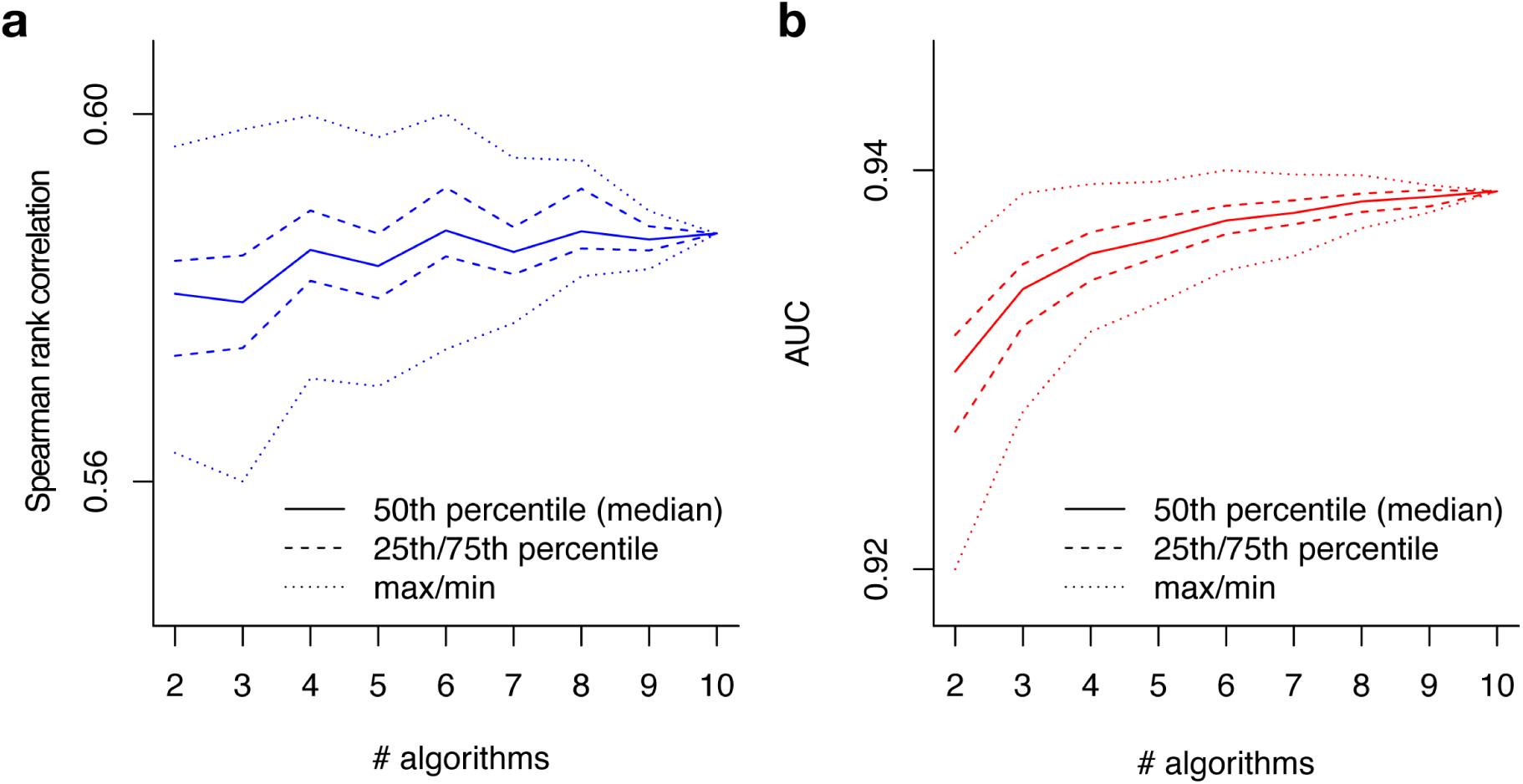
Performance of ensembles. This figure shows the performance in the (**a**) correlation metric and (**b**) AUC metric of the ensemble based on the median prediction of all possible combinations of a given size of the top 9 algorithms plus the benchmark. The median performance of the ensemble tends to increase with its size. However, the maximum performance in both metrics tends to plateau (or even decrease) after the ensemble reaches a size equal to 3.

## 4 Discussion

We created a novel dataset of L1000 profiles for over 120 shRNA and compound experiments with several replicates for a total of 2,200 gene expression profiles of genes measured independently, and in tandem. This dataset constitutes now a public resource (S1 Data) to all the researchers in this area who are interested in testing their deconvolution approaches.

Using an open innovation competition, we collected and evaluated multiple and diverse deconvolution methods. The best approach was based on a random forest, which is a collection of decision tree regressors. This method achieved: (i) the highest global correlation between the ground-truth and the corresponding deconvoluted data, (ii) the lowest interreplicate variation, and (iii), compared to the benchmark, was able to detect more than a thousand additional extremely modulated genes, while reducing the false positives at the same time. Our analysis further showed that these gains are considerable when the gene populations were sampled in different proportions (here, genes in high and low bead proportions), with the k-means benchmark approach being systematically less accurate because it does little to mitigate the discrepancy in variability between the genes measured with high and low bead proportion.

In addition, the random-forest approach achieved these improvements with only 10 trees on 60 features. Thus, the algorithm is also relatively fast and easy to implement. By comparison, the fastest approach used a more traditional Gaussian mixture model (with plate-level adjustments), which turned out to be less accurate. Hence, and overall, our analysis provided evidence of the tremendous potential of using random-forest methods for deconvolution problems in biology.

## 5 Acknowledgements

This work was funded by the Eric and Wendy Schmidt Foundation, NASA Center of Collaborative Excellence, and the Kraft Precision Medicine Accelerator & Division of Research and Faculty Development at the Harvard Business School. The CMap competitions were supported in part by the NIH Common Funds Library of Integrated Network-based Cellular Signatures (LINCS) program U54HG006093 and NIH Big Data to Knowledge (BD2K) program 5U01HG008699.

## Supporting information

**S1 Table.**
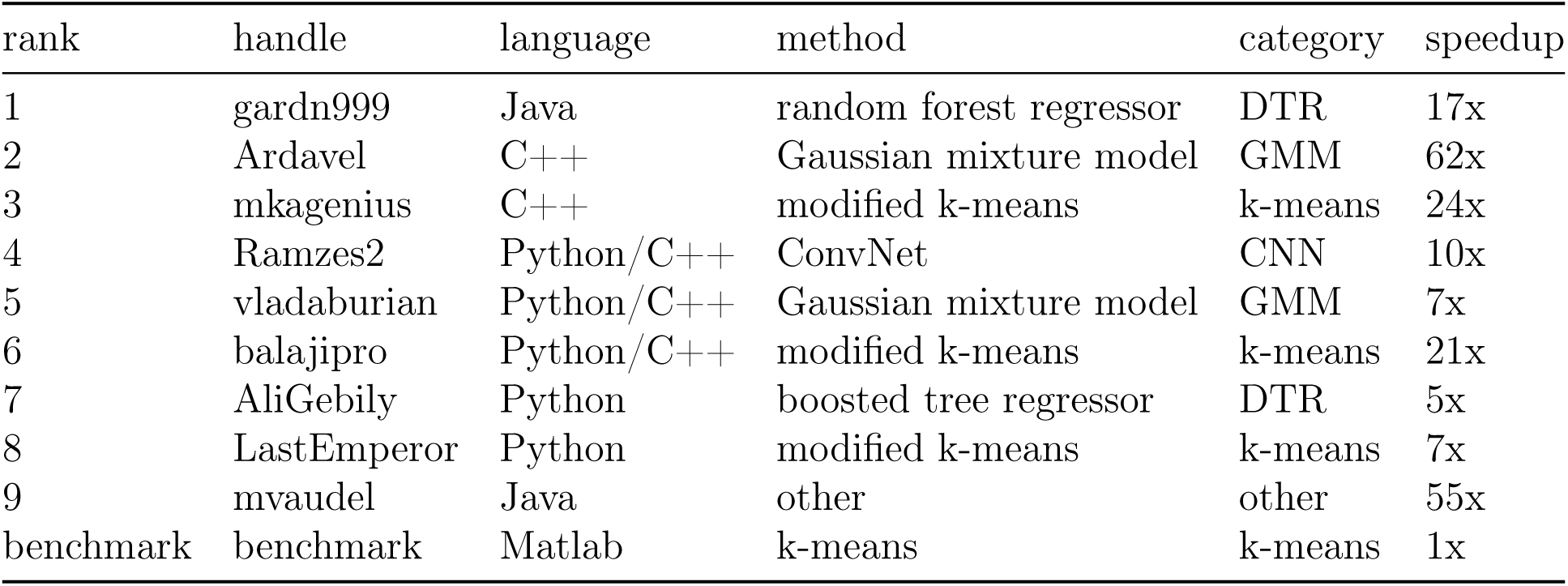
Top-nine performing solutions and benchmark. This table lists the top-nine solutions and the languages and algorithms each used, as well as the average speedup per plate relative to the k-means benchmark.

**S2 Table.**
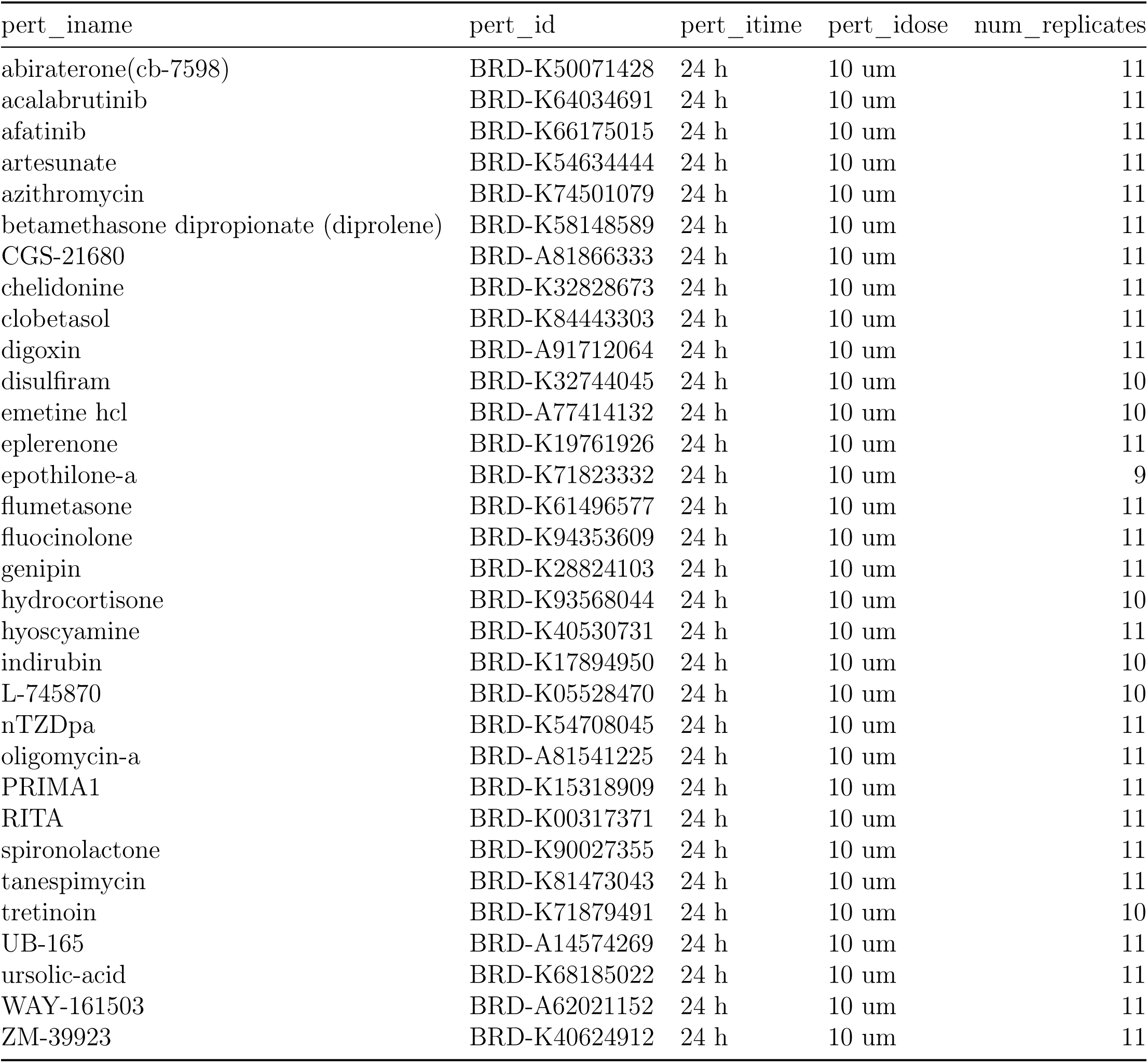
Compound perturbagens descriptives. This table shows componud perturbagen names (`pert_iname`), unique id (`pert_id`), time of treatment (`pert_itime`), dose (`pert_idose`), and number of replicates (`num_replicates`).

**S3 Table.** Short-hairpin (shRNA) perturbagens descriptives. This table shows shRNA perturbagen names (`pert_iname`), unique id (`pert_id`), and number of replicates (`num_replicates`).

| pert_iname | pert_id | num_replicates |
| --- | --- | --- |
| ABCB6 | TRCN0000060320 | 4 |
| ADI1 | TRCN0000052275 | 4 |
| ALDOA | TRCN0000052504 | 4 |
| ANXA7 | TRCN0000056304 | 4 |
| ARHGAP1 | TRCN0000307776 | 4 |
| ASAH1 | TRCN0000029402 | 4 |
| ATMIN | TRCN0000141397 | 4 |
| ATP2C1 | TRCN0000043279 | 4 |
| B3GNT1 | TRCN0000035909 | 4 |
| BAX | TRCN0000033471 | 4 |
| BIRC5 | TRCN0000073718 | 4 |
| BLCAP | TRCN0000161355 | 4 |
| BLVRA | TRCN0000046391 | 4 |
| BNIP3L | TRCN0000007847 | 4 |
| CALU | TRCN0000053792 | 4 |
| CCDC85B | TRCN0000242754 | 4 |
| CCND1 | TRCN0000040038 | 4 |
| CD97 | TRCN0000008234 | 4 |
| CHMP4A | TRCN0000150154 | 4 |
| CNOT4 | TRCN0000015216 | 4 |
| DDR1 | TRCN0000000618 | 4 |
| DDX10 | TRCN0000218747 | 4 |
| DECR1 | TRCN0000046516 | 4 |
| DNM1L | TRCN0000001097 | 3 |
| ECH1 | TRCN0000052455 | 4 |
| EIF4EBP1 | TRCN0000040206 | 4 |
| EMPTY_VECTOR | TRCN0000208001 | 15 |
| ETFB | TRCN0000064432 | 4 |
| FDFT1 | TRCN0000036327 | 4 |
| GALE | TRCN0000049461 | 4 |
| GFP | TRCN0000072181 | 16 |
| GRN | TRCN0000115978 | 4 |
| GTPBP8 | TRCN0000343727 | 4 |
| HDGFRP3 | TRCN0000107348 | 4 |
| HIST1H2BK | TRCN0000106710 | 4 |
| IKBKAP | TRCN0000037871 | 4 |
| INPP4B | TRCN0000230838 | 4 |
| INSIG1 | TRCN0000134159 | 4 |
| ITFG1 | TRCN0000343702 | 3 |
| JMJD6 | TRCN0000063340 | 4 |
| LBR | TRCN0000060460 | 4 |
| LGMN | TRCN0000029255 | 4 |
| LPGAT1 | TRCN0000116066 | 4 |
| LSM6 | TRCN0000074719 | 4 |
| MAPKAPK2 | TRCN0000002285 | 4 |
| MAPKAPK3 | TRCN0000006154 | 4 |
| MAPKAPK5 | TRCN0000000684 | 4 |
| MIF | TRCN0000056818 | 4 |
| MRPL12 | TRCN0000072655 | 4 |
| NT5DC2 | TRCN0000350758 | 4 |
| NUP88 | TRCN0000145079 | 4 |
| PARP2 | TRCN0000007933 | 4 |
| PLCB3 | TRCN0000000431 | 4 |
| POLE2 | TRCN0000233181 | 4 |
| PPIE | TRCN0000049371 | 4 |
| PRKAG2 | TRCN0000003146 | 4 |
| PSMB10 | TRCN0000010833 | 4 |
| PTPN6 | TRCN0000011052 | 4 |
| RAB11FIP2 | TRCN0000322640 | 4 |
| RALB | TRCN0000072956 | 4 |
| RHEB | TRCN0000010425 | 3 |
| RNF167 | TRCN0000004100 | 4 |
| RPN1 | TRCN0000072588 | 4 |
| SLC25A4 | TRCN0000044967 | 4 |
| SNX11 | TRCN0000127684 | 4 |
| STK25 | TRCN0000006270 | 4 |
| STUB1 | TRCN0000007525 | 4 |
| STXBP1 | TRCN0000147480 | 4 |
| SYPL1 | TRCN0000059926 | 4 |
| TATDN2 | TRCN0000049828 | 4 |
| TM9SF3 | TRCN0000059371 | 4 |
| TMEM110 | TRCN0000127960 | 4 |
| TMEM50A | TRCN0000129223 | 4 |
| trcn0000014632 | TRCN0000014632 | 4 |
| trcn0000040123 | TRCN0000040123 | 4 |
| trcn0000220641 | TRCN0000220641 | 4 |
| trcn0000221408 | TRCN0000221408 | 4 |
| trcn0000221644 | TRCN0000221644 | 4 |
| TSKU | TRCN0000005222 | 4 |
| UGDH | TRCN0000028108 | 4 |
| USP14 | TRCN0000007428 | 4 |
| USP6NL | TRCN0000253832 | 4 |
| VAT1 | TRCN0000038193 | 4 |
| VDAC1 | TRCN0000029126 | 4 |
| WIPF2 | TRCN0000029833 | 4 |
| YME1L1 | TRCN0000073864 | 4 |
| ZW10 | TRCN0000155335 | 4 |

### S1 Data

LINK TO DATA REPOSITORY on http://CLUE.io WILL BECOME AVAILABLE AFTER PUBLICATION

## S1 Appendix Scoring function

This appendix describes the scoring function used in the contest to evaluate the performance of the competitors’ submissions.

Submissions were scored based on a scoring function that combines measures of accuracy and computational speed. Accuracy measures were obtained by comparing the contestant’s predictions, which were derived from *DUO* data, to the equivalent *UNI* ground truth data generated from the same samples.

The scoring function combines two measures of accuracy: correlation and AUC, which are applied to deconvoluted (*DECONV*) data and one to differential expression (*DE*) data, respectively.

*DE* is derived from DECONV by applying a series of transformations (parametric scaling, quantile normalization, and robust z-scoring) that are described in detail in Subramanian et al. ^1^. The motivation for scoring *DE* data in addition to *DECONV* is because it is at this level where the most biologically interesting gene expression changes are observed. Of particular interest is obtaining significant improvement in the detection of, so called, “extreme modulations.” These are genes that notably up- or down-regulated by perturbation and hence exhibit an exceedingly high (or low) *DE* values relative to a fixed threshold.

The first accuracy component is based on the Spearman rank correlation between the predicted *DECONV* data and the corresponding *UNI* ground truth data.

For a given dataset *p*, let *M*DUO*,p* and *M*UNI*,p* denote the matrices of the estimated gene intensities for *G* = 976 genes (rows) and *S* = 384 experiments (columns) under DUO and UNI detection. Compute the Spearman rank correlation matrix, *ρ*, between the rows of these matrices and take the median of the diagonal elements of the resulting matrix (i.e., the values corresponding to the matched experiments between UNI and DUO) to compute the median correlation per dataset,

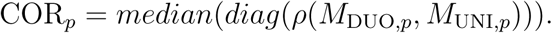

The second component of the scoring function is based on the Area Under the receiver operating characteristic Curve (AUC) that uses the competitor’s DE values at various thresholds to predict the UNI’s DE values being higher than 2 (“high”) or lower than −2 (“low”).

For a given dataset *p*, let AUC*p,c* denote the corresponding area under the curve where *c* = *{*high, low*}*; then, compute the arithmetic mean of the area under the curve per class to obtain the corresponding score per dataset:

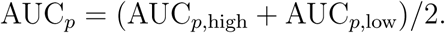

These accuracy components were integrated into a single aggregate scores:

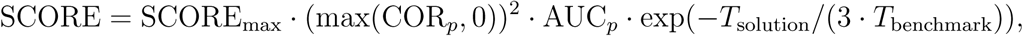

where *T*solution is the run time for deconvoluting the data in each plate, and *T*benchmark is the deconvolution time required by the benchmark dpeak implementation.

## S2 Appendix L1000 Experimental Scheme

The L1000 assay uses Luminex bead-based fluorescent scanners to detect gene expression changes resulting from treating cultured human cells with chemical or genetic perturbations [Subramanian 2017]. Experiments are performed in 384-well plate format, where each well contains an independent sample. The Luminex scanner is able to distinguish between 500 different bead types, or colors, which CMap uses to measure the expression levels of 978 landmark genes using two detection approaches.

In the first detection mode, called *UNI*, each of the 978 landmark genes is measured individually on one of the 500 Luminex bead colors. In order to capture all 978 genes, two detection plates are used, each measuring 489 landmarks. The two detection plates’ worth of data are then computationally combined to reconstruct the full 978-gene expression profile for each sample.

By contrast, in the *DUO* detection scheme two genes are measured using the same bead color. Each bead color produces an intensity histogram which characterizes the expression of the two distinct genes. In the ideal case, each histogram consists of two peaks each corresponding to a single gene. The genes are mixed in a 2:1 ratio, thus the areas under the peaks have a 2:1 ratio (see Figure 1), which enables the association of each peak with the specific gene. The practical advantage of the DUO detection mode is that it uses half of the laboratory reagents as UNI mode, and hence *DUO* is and has been the main detection mode used by CMap. After *DUO* detection, the expression values of the two genes are computationally extracted in a process called ‘peak deconvolution.’ See Subramanian et al. ^1^ for more details.

## Footnotes

1 These results are for the dataset with shRNA experiments for which we know the targeted genes. We expect similar results for the dataset of compound experiments.

## References

[1] Aravind Subramanian, Rajiv Narayan, Steven M Corsello, David D Peck, Ted E Natoli, Xiaodong Lu, Joshua Gould, John F Davis, Andrew A Tubelli, Jacob K Asiedu, et al. A next generation connectivity map: L1000 platform and the first 1,000,000 profiles. Cell, 171(6):1437–1452, 2017.

[2] Brian Cleary, Le Cong, Anthea Cheung, Eric S Lander, and Aviv Regev. Efficient generation of transcriptomic profiles by random composite measurements. Cell, 171(6): 1424–1436, 2017.

[3] Shai S Shen-Orr, Robert Tibshirani, Purvesh Khatri, Dale L Bodian, Frank Staedtler, Nicholas M Perry, Trevor Hastie, Minnie M Sarwal, Mark M Davis, and Atul J Butte. Cell type–specific gene expression differences in complex tissues. Nature methods, 7(4): 287, 2010.

[4] Yi Zhong and Zhandong Liu. Gene expression deconvolution in linear space. Nature methods, 9(1):8, 2012.

[5] Aaron M Newman, Chih Long Liu, Michael R Green, Andrew J Gentles, Weiguo Feng, Yue Xu, Chuong D Hoang, Maximilian Diehn, and Ash A Alizadeh. Robust enumeration of cell subsets from tissue expression profiles. Nature methods, 12(5):453, 2015.

[6] Konstantin Zaitsev, Monika Bambouskova, Amanda Swain, and Maxim N Artyomov. Complete deconvolution of cellular mixtures based on linearity of transcriptional signatures. Nature communications, 10(1):2209, 2019.

[7] Yue Deng, Feng Bao, Qionghai Dai, Lani F Wu, and Steven J Altschuler. Scalable analysis of cell-type composition from single-cell transcriptomics using deep recurrent learning. Nature methods, 16(4):311, 2019.

[8] Peng Lu, Aleksey Nakorchevskiy, and Edward M Marcotte. Expression deconvolution: a reinterpretation of dna microarray data reveals dynamic changes in cell populations. Proceedings of the National Academy of Sciences, 100(18):10370–10375, 2003.

[9] Stephan Preibisch, Fernando Amat, Evangelia Stamataki, Mihail Sarov, Robert H Singer, Eugene Myers, and Pavel Tomancak. Efficient bayesian-based multiview deconvolution. Nature methods, 11(6):645, 2014.

[10] Shai S Shen-Orr and Renaud Gaujoux. Computational deconvolution: extracting cell type-specific information from heterogeneous samples. Current opinion in immunology, 25(5):571–578, 2013.

[11] Karim R Lakhani, Kevin J Boudreau, Po-Ru Loh, Lars Backstrom, Carliss Baldwin, Eric Lonstein, Mike Lydon, Alan MacCormack, Ramy A Arnaout, and Eva C Guinan. Prize-based contests can provide solutions to computational biology problems. Nature biotechnology, 31(2):108, 2013.

[12] Benjamin M Good and Andrew I Su. Crowdsourcing for bioinformatics. Bioinformatics, 29(16):1925–1933, 2013.

[13] Andrea Blasco, Michael G. Endres, Rinat A. Sergeev, Anup Jonchhe, N. J. Maximilian Macaluso, Rajiv Narayan, Ted Natoli, Jin H. Paik, Bryan Briney, Chunlei Wu, Andrew I. Su, Aravind Subramanian, and Karim R. Lakhani. Advancing computational biology and bioinformatics research through open innovation competitions. PLOS ONE, 14(9): 1–17, 09 2019. doi: 10.1371/journal.pone.0222165. URL https://doi.org/10.1371/journal.pone.0222165.

[14] Olaf Ronneberger, Philipp Fischer, and Thomas Brox. U-net: Convolutional networks for biomedical image segmentation. In International Conference on Medical image computing and computer-assisted intervention, pages 234–241. Springer, 2015.

[15] Laurens van der Maaten and Geoffrey Hinton. Visualizing data using t-sne. Journal of machine learning research, 9(Nov):2579–2605, 2008.

